# Mechanistic Insights into ASO-RNA Complexation: Advancing Antisense Oligonucleotide Design Strategies

**DOI:** 10.1101/2024.04.18.590021

**Authors:** Johanna Hörberg, Antonio Carlesso, Anna Reymer

## Abstract

Oligonucleotide drugs, an emerging modulator class, hold promise for targeting previously undruggable biomacromolecules. To date, only 18 oligonucleotide drugs, including sought-after antisense oligonucleotides (ASO) and splice-switching oligonucleotides (SSO), have FDA approval. These agents effectively bind mRNA, inducing degradation or modulating splicing. Current oligonucleotide drug design strategies prioritize full Watson-Crick base pair complementarity, overlooking mRNA target 3D shapes. Given that mRNA conformational diversity can impact hybridization, incorporating mRNA key-structural properties into the design may expedite ASO lead discovery. Using atomistic molecular dynamics simulations inspired by experimental data, we demonstrate the advantages of incorporating common triple base pairs into the design of antisense oligonucleotides (ASOs) targeting RNA hairpin motifs, which are highly accessible regions for interactions. By employing an RNA pseudoknot modified into an ASO-hairpin complex, we investigate the effects of ASO length and hairpin loop mutations. Our findings suggest that ASO-mRNA complex stability is influenced by ASO length, number of common triple base pairs, and the dynamic accessibility of bases in the hairpin loop. Our study offers new mechanistic insights into ASO-mRNA complexation and underscores the value of pseudoknots in constructing training datasets for machine learning models aimed at designing novel ASO leads.

## Introduction

Since 1990s, oligonucleotide therapeutics emerge as a promising class of drugs offering the attractive potential to target any genetic target, including the 10,000 proteins previously classified as undruggable by conventional small molecules.(1, 2) Despite their potential, only 18 oligonucleotide drugs have been approved by FDA, where the half constitutes antisense oligonucleotides (ASOs) (Table S1).(3) The first FDA-approved ASO, fomivirsen, appeared in 1998 for the treatment of cytomegalovirus retinitis. (3, 4) Thereafter, additional eight ASOs have been approved, targeting rare genetic and neurodegenerative disorders. (3) To this date, there are still no oligonucleotide therapeutics approved for treatment of cancer; though there are ongoing clinical trials, but most of them are still in phase I-II.(5, 6)

ASOs are short stretches of 12-30 nucleotides single-stranded nucleic acid sequences (ssDNA or ssRNA) that bind target RNA, most commonly messenger-RNA (mRNA) through Watson-Crick base pairing.(5, 7) By binding to their target, ASOs control protein synthesis through different mechanisms: (i) enhance degradation through recruitment of endonucleases like RNAse H, (ii) act as steric blocks or (iii) modulate the splicing, a class also known as splice-switching oligonucleotides (SSO).(2, 3, 5, 7) Despite their considerable potential and significant investment into their development, antisense therapeutics, like conventional small drugs, encounter clinical trial setbacks related to toxicity, potency, efficacy, and specificity.(2) In addition, ASOs are susceptible to degradation by nucleases, and must contain several modifications in their backbones (5, 7–9), which often lead to unwanted off-target effects.(10, 11). Thus, to advance ASO therapeutics, we must deepen our understanding of the mechanistic principles governing the relationship between ASO sequence design and target specificity.(12)

Current ASOs design strategies follow the “full Watson-Crick base pairing principle”, where an ASO strand is designed to be complementary to a selected mRNA region. Since a 12-30-mer oligonucleotide sequence is likely to occur only once,(3) the antisense approach anticipates achieving high selectivity through only the knowledge of the mRNA target sequence.(2, 3) However, firstly, RNA base pairing can tolerate mismatches.(13) Secondly, because of the hydrophobicity of nucleobases and the backbone flexibility, mRNA can fold into different 3D motifs which can interfere with the mRNA-ASO hybridization and the selectivity.(12) Thus, one either tries to target more unstructured regions of mRNA(14) or screen large libraries of different ASOs.(2, 12)

Recent studies have highlighted the importance of taking the 3D structure of RNA targets into account in the design of ASOs. Lulla et al.(15) used structural information (icSHAPE and X-ray) of the conserved 3’ stem-loop II-like motif (s2m) in the 3’-untranslational region of the SARS-CoV2. Researchers obtained high affinity ASOs by targeting regions with computationally predicted exposed bases of s2m, which could facilitate the initial ASO-target mRNA interactions. SHAPE experiments revealed changes in the s2m region induced by the ASO binding, triggering an efficient RNAse H cleavage. The designed ASOs exhibited low toxicity and were able to inhibit the replication of both SARS-CoV2 and distantly related RNA viruses, astroviruses, in a dose-dependent manner.

In another study, Li et al.(12) proposed a structure-based method for ASO design, demonstrating the advantage of ASOs being compatible with target RNA structures in three-dimensional (3D) space through the formation of common triple base pairs (b.p.) and Hogsteen b.p. interactions. The researchers used the human telomerase RNA pseudoknot as a model system, which they modified into an ASO-hairpin complex. By performing isothermal titration calorimetry experiments, Li et al. showed that truncation of the ASO and mutations of the target-RNA hairpin decreased the affinity by 20 and 130-fold, respectively. The authors proposed that the decrease in affinity is attributed to a reduction in interactions involving common triple base pairs. Building on available information on favorable triple and Hogsteen b.p. interactions, Li et al. designed ASOs targeting the frameshift stimulation element and transcription regulatory sequence of SARS-CoV-2, successfully inhibiting the viral replication in human cells.

Inspired by the study of Li et al.(12), we further explore the mechanistic details of ASO-hairpin complexation. Utilizing the human telomerase RNA pseudoknot(16), modified into an ASO-hairpin complex, we conduct microsecond-long all-atomistic molecular dynamics simulations for the unmodified, ASO truncated, and mutated RNA hairpin loop variants (Figure 1). Through detailed analyses of the dynamics and interactions within the ASO-RNA complexes, our data highlight two critical factors at the molecular level: Firstly, the disruption of common triple base pairs within the ASO-hairpin complex; and secondly, the dicreased accessibility of bases within the hairpin loop can contribute significantly to the observed elevated dissociation rates.

**Figure 1:**
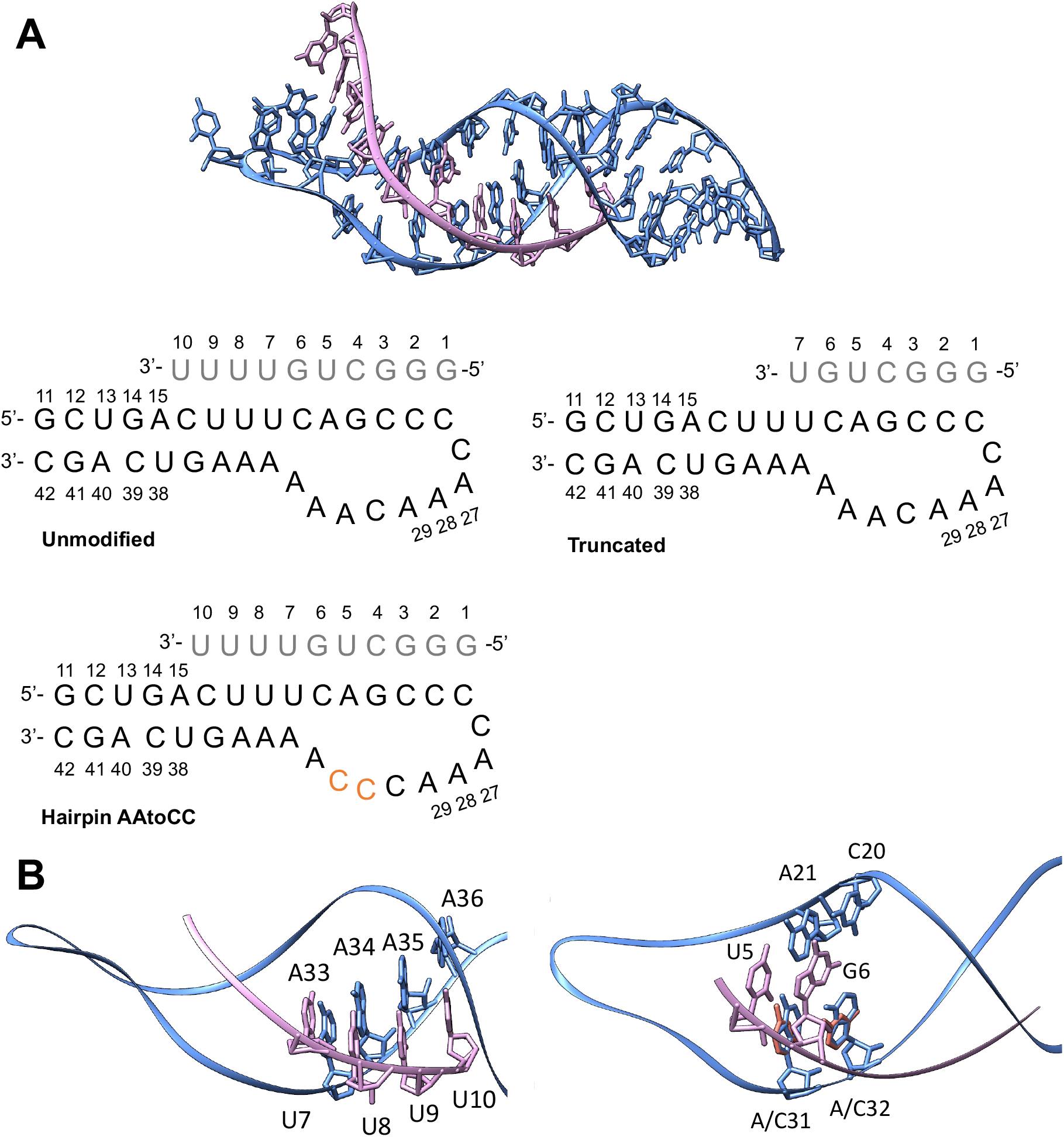
**A**. ASO-hairpin model systems prepared from the NMR structure of human RNA telomerase pseudoknot (PDB: 1YMO).(16) **B**. Left-hand panel: ASO “tail” interactions with the RNA hairpin stem, where the interactions between U8-U10 and A34-A36 are lost upon the ASO truncation. Right-hand panel: triple b.p. interactions involving the ASO nucleotides U5 and G6. These are suggested to brake upon A31A32 ->C31C32 (orange color, panel A) mutations.

Furthermore, our investigation reveals that the accessible bases within the hairpin loop not only serve as interaction sites for ASOs but also plays a crucial role in shaping a suitable binding pocket to accommodate the ASO. Our mechanistic data underscores the dependence of ASO-RNA complex stability on both the length of the ASO and the abundance of common triple base pairs. Our computational study highlights the potential utility of pseudoknots and triple helix complexes as valuable models for generating training datasets, which can advance the development of machine learning tools for designing novel ASO leads. We also emphasize the importance of integrating structural information for mRNA targets, offering significant potential to streamline the ASO design process by directing attention toward mRNA regions readily accessible for potential 3D interactions with ASOs.

## Methods

### Systems Preparation

We prepare three ASO-target RNA hairpin systems (Figure 1A): later referred to as “unmodified”, “truncated” ASO and the target RNA hairpin mutant “AAtoCC”, from the NMR structure of the human RNA telomerase pseudoknot (PDB ID: 1YMO)(16), using USCF Chimera(17). The NMR ensemble contains 20 structures, where we select the first snapshot as the model system. We remove four nucleotides, connecting the ASO to the hairpin, to obtain the unmodified system. For the truncated ASO system, we remove the nucleotides 8-10 of the ASO. For the AAtoCC system, we exchange the nucleotides A31 and A32 of the target-RNA hairpin with cytosines.

### Molecular Dynamics Simulations

We perform all molecular dynamics simulations with the GROMACS MD engine version 2021.6(18), using the parmbsc0+chiOL3(19, 20) forcefield. Each system is solvated with 15Å TIP3P water(21) in cubic boxes. Charges are neutralised with Mg2+. Additional 150mM NaCl is added to reach a physiological salt concentration.(22) For RNA, a Mg2+ and Na+ environment is preferred as it provides the greater structural stability.(22–24) Each system is subjected to energy minimization with 5000 steps of steepest decent, followed by 500 ps and 20 ns equilibration-runs with weak position restraints on heavy atoms of the solute (1000 kJ/mol) in the NVT and NPT ensembles respectively, adjusting temperature and pressure to 300 K and 1 atm,(25, 26) under periodic boundary conditions. Releasing the restraints, for each of the systems, for each system we carry out 2000ns MD simulations at constant pressure and temperature (1 atm and 300 K).

### Molecular Dynamics Trajectories Analysis

The trajectories are analysed using both GROMACS analyses tools(18) and CPPTRAJ program from AMBERTOOLS 16 software package.(27) RMSD, RMSF and radius of gyration are analysed for heavy atoms using GROMACS. We analyse nucleic acids contacts using the CPPTRAJ. We distinguish between specific (hydrogen bonding and hydrophobic contacts formed between atoms of the nucleobases) and nonspecific contacts (that involved interactions with at least one backbone atom). For every pair of ASO-target-RNA hairpin residues or hairpin-hairpin residues, we sum up all hydrogen bonds, salt bridges, and hydrophobic (apolar) interactions, where we set the contribution of each type of contact to 1. This approximation is done for simplicity, since the energy cost of hydrogen bonds, salt bridges, and hydrophobic interactions varies greatly, depending on the nature of the atoms involved, the bond geometry and the surrounding environment. Only the contacts that were present for longer than 10% of the trajectory are considered, see previous publications for further details.(28, 29) The GROMACS energy tool is used to calculate the ASO-target-RNA hairpin interaction energies, which operates with the GROMACS force field functional terms i.e., the short range (within a cut-off distance ∼10Å) electrostatic (Coulomb) and Lennard-Jones (vdW) interactions for every atom pair within the selected groups are calculated and summed up to obtain the total interaction energy. We separate interaction energies into specific and nonspecific,(30) by calculating interactions for four atom groups: 1. between ASO and target-RNA hairpin nucleobases (ASOBase-hairpinBase). 2. Between ASO and target-RNA hairpin backbone (ASOBac-hairpinBac). 3. Between ASO nucleobases and target-RNA hairpin backbone (ASOBase-hairpinBac). 4. Between ASO backbone and target-RNA hairpin nucleobases (ASOBac-hairpinBase). The first group includes the specific interactions, whereas the remaining three groups constitute the nonspecific interactions. We also perform free energy analysis and entropy analysis, using the MMPBSA/MMGBSA method (31, 32) included in the AMBER CPPTRAJ module. We use the GROMACS “covar” and “anaeig” tools for the principal component analysis (PCA) of the ASO-target RNA complex heavy atoms.

## Additional Information

We use MATLAB for the generation of all plots and USCF Chimera(17) for molecular graphics.

## Results

In this study, we aimed to elucidate the mechanistic principles governing the stable and selective binding of ASOs to their RNA targets and to lay the foundation for future CAD routines for the rational design of ASOs. We drew inspiration from Li et al. study(12) that, based on *in vitro* binding experiments, proposed that an ASO-target-RNA complex forms through common triple and Hoogsteen base pairing. Using an ASO-hairpin complex (Figure 1A), derived from the human RNA telomerase pseudoknot (PDB ID: 1YMO)(16), Li et al. demonstrated that truncating the ASO or mutating the RNA hairpin loop (A31A32->C31C32) decreased the affinity by 20- and 130-fold, respectively. Both alterations were attributed to a reduction in the number of triple base pair interactions. Specifically, the truncated ASO variant was suggested to result in the loss of interactions with the three adenines (A34-A36) of the hairpin stem, while the AA to CC hairpin-mutated variant led to non-compatible triple base pairs interactions for U5 and G6 (Figure 1B). These mechanistic conclusions were drawn solely from the analysis of the statically solved structure of the pseudoknot (PDB ID: 1YMO)(16). However, nucleic acids are extremely dynamic molecules. This motivated us to conduct atomistic molecular dynamics (MD) simulations to rationalize the experimental data and derive additional mechanistic insights into the importance of triple b. p. interactions for ASO-RNA hairpin complexation. We aimed to explain at a molecular level why the loss of interactions with the hairpin loop may have a more significant impact on the dissociation rate than the interactions with the stem. Additionally, we seeked to identify other potential structure-mechanistic aspects that could contribute to the reduced affinity of the ASO-hairpin complex.

We prepared three ASO-RNA hairpin systems (unmodified, truncated, AAtoCC) from the RNA telomerase pseudoknot using UCSF Chimera and subjected each complex to two microseconds of atomistic MD simulations. Following the simulations, we analyzed various mechanistic and thermodynamic parameters. We started by assessing the root mean square deviation (RMSD) of heavy atoms within the complex, the RNA hairpin, and the ASO, along with the radius of gyration along the trajectories (see Figure 2A).

**Figure 2:**
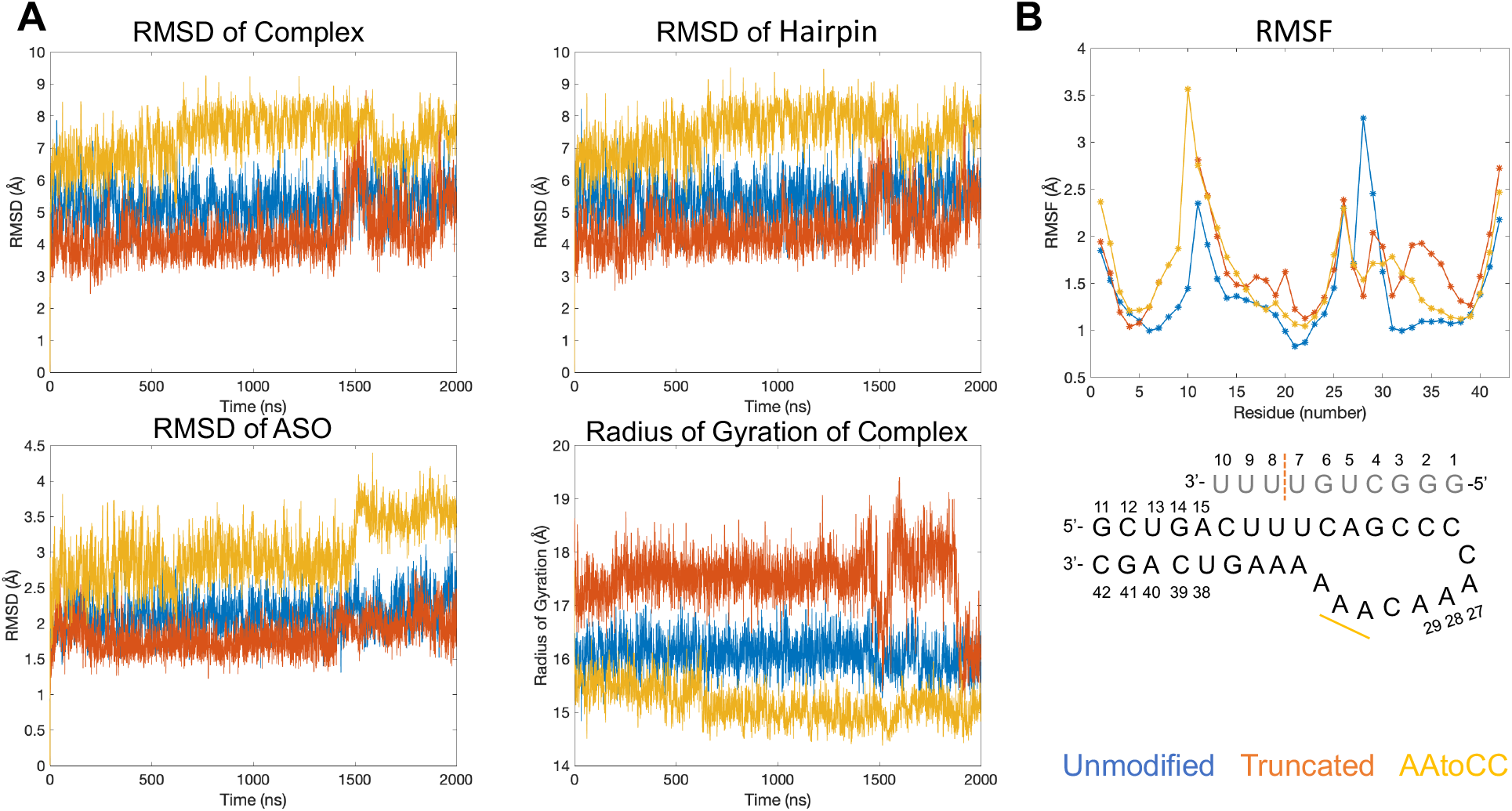
**A**. Evolution of the RMSD and radius of gyration values of the heavy atoms relative to the starting structure. **B**. Per residue RMSF for the MD trajectory window 100-2000ns for the unmodified and AAtoCC systems and 100-1400ns for the truncated ASO system. In the schematic figure of the ASO-hairpin, the truncation is marked with orange dashed line and the mutations are marked with yellow bold line.

The choice of the initial structure as the reference state allowed us to compare changes resulting from the ASO truncation and the RNA hairpin loop mutations. The AAtoCC system exhibited the most substantial deviation from the initial structure, mainly due to structural changes in the RNA hairpin loop. Additionally, the ASO showed increased flexibility, marked by a significant rise in RMSD after 1500 ns, as the complex transitioned into a more compact state with a lower radius of gyration. The truncated ASO system demonstrated stable RMSD fluctuations until the final 500 ns, which coincided with significant changes in the radius of gyration, when the complex became more compact. In contrast, the unmodified system displayed similar RMSD and radius of gyration fluctuations along the entire trajectory, suggesting a more stable complex, though also with a substantial deviation from the initial structure. Deviation from the initial structures is expected since the complexes were derived from the telomerase pseudoknot, which undergoes conformational changes upon the transformation into ASO-hairpin complexes. The pseudoknot’s “ASO”-arm contains an additional four nucleotides connecting to the 5’-terminal of the “RNA hairpin” resulting in a more rigid system. Further truncation of the ASO (as is in the truncated system), removes contacts with the hairpin stem increasing the conformational freedom for the RNA hairpin, leading to larger fluctuations in the radius of gyration (see Figures 2A and S1).

To gain further insights into the dynamic differences among the three systems, we analyzed the Root Mean Square Fluctuation (RMSF) per residue (Figure 2B), disregarding the first 100 ns as equilibration. In the case of the truncated ASO system, we excluded also the last 600 ns due to significant RMSD fluctuations, which resulted in incompatible RMSF values compared to the other systems (see Figure S2). As illustrated in Figure 2B, the increased fluctuations of the ASO in the AAtoCC system can be attributed to greater flexibility in the ASO tail (bases 7-10). Similarly, the truncated ASO system also displayed enhanced flexibility in the terminal ASO bases (bases 6-7) compared to the unmodified system. Additionally, both the truncated ASO and the AAtoCC systems exhibited increased flexibility in the hairpin loop region when compared to the unmodified system, except for the A28-A29 bases, which aligns with a naturally more flexible region observed in the NMR structure of the telomerase pseudoknot. Higher RMSF values suggest a more dynamic unsettled complex.

In conclusion, based on the analyses of RMSD, RMSF, and radius of gyration, our simulations follow the trend of the experimentally determined dissociation constants, ranking AAtoCC > truncated ASO > unmodified.

### Interaction Energies and Free Energy Calculations

We proceed with the analyses of the thermodynamic parameters of the ASO-RNA hairpin complexes. It must be noted that *in silico* estimation of thermodynamic parameters, such as interaction energies or binding free energies for nucleic acid complexes constitutes a challenge due to the high negative charge of the molecular backbones. Nevertheless, to see if available computational methods can qualitatively capture the binding experiments trend, we next derive the interaction energies, using both the GROMACS energy tool and the MMGBSA/MMPBSA method.

The GROMACS energies include the short-range electrostatic (Columb) and vdW-(Lennard Jones) interactions and are derived from the force field nonbonded interaction components. We derive the interaction energies for four groups: 1) ASO and hairpin nucleobases atoms (ASOBase-hairpinBase), 2) ASO and hairpin backbones atoms (ASOBac-hairpinBac), 3) ASO nucleobases and hairpin backbone atoms (ASOBase-hairpinBac), and 4) ASO backbone and hairpin nucleobases atoms (ASOBac-hairpinBase). The interactions in the first group, we call the specific interactions, whereas the interactions in the remaining three groups are referred to as nonspecific interactions. We also combine the four groups to calculate the total interactions energies (Figures S3-S4, Table 1).

**Table 1:** GROMACS interaction energies between various atom groups of the ASO-hairpin complexes compared to the dissociation constants derived from ITC experiments.(12)

|  | ASOBase-hairpinBase<br>(kcal/mol) | ASOBac-hairpinBac<br>(kcal/mol) | ASOBase-hairpinBac<br>(kcal/mol) | ASOBac-hairpinBase<br>(kcal/mol) | Total<br>(kcal/mol) | Kd<br>(nM) |
| --- | --- | --- | --- | --- | --- | --- |
| Unmod | -223.4±7.2 | -14.2±4.5 | -14.0±3.2 | -53.1±7.6 | -304.7±15.6 | 28±4 |
| Trunc | -168.6±12.3 | -4.0±2.0 | -11.0±3.2 | -44.6±5.2 | -228.2±13.1 | 590±160 |
| AAtoCC | -222.9±9.6 | -16.8±4.6 | -8.2±3.2 | -52.1±7.6 | -299.9±14.3 | 3600±1600 |

As illustrated in Table 1, specific interactions contribute the most to the total interaction energy, given that interactions with backbone atoms may lead to some repulsion. Based on the GROMACS interaction energies, we cannot reproduce the trend of the experimental dissociation constants.(12) The truncated ASO system shows the least favorable interaction energies, while the unmodified and AAtoCC systems display similar interaction energies that overlap within standard deviations. This is because, the GROMACS interactions energies method depends on the number of contacts. Thus, since the ASO truncation removes three bases and consequently several specific interactions, the interaction energies will be noticeably reduced. In addition, the method cannot accurately discriminate between interactions formed with an adenine NH2-group versus a cysteine NH2-group, whereas nature might favor one type of contact over the other. Nevertheless, some interesting insights can be derived from the fluctuations of the interaction energies along the trajectories (Figure S3). For the specific interactions, both the truncated ASO and the AAtoCC systems exhibit large fluctuations towards the end of the trajectory, which correlate with the RMSD analysis, and may suggest the beginning of ASO-hairpin dissociation. The unmodified system exhibits a reduction in nonspecific interactions towards the end of the trajectory, suggesting a conformational change (see discussion about the contacts networks below). The progression of the total energy (Figure S4) shows more fluctuations for both the truncated ASO and the AAtoCC systems, whereas the unmodified system exhibits two modes, one at -310 kcal/mol until 1500 ns and thereafter a reduction to -284 kcal/mol.

The binding free energies calculated using the MMPBSA/MMGBSA method do not align with the trend of the experimental dissociation constants(12) either (Table 2). Firstly, the truncated ASO system is estimated to be the most favorable complex and the AAtoCC system – the least. This is because, the electrostatic terms (PBELE and GBELE) are positive, i.e., repulsive. However, for the truncated system where the ASO lacks the three last bases, the electrostatic interactions become less repulsive. Conversely, the AAtoCC system exhibits the most repulsive electrostatics. This is in line with the radius of gyration results, where the more compact complex may lead to shorter phosphate-phosphate distances compared to the other systems.

**Table 2:**
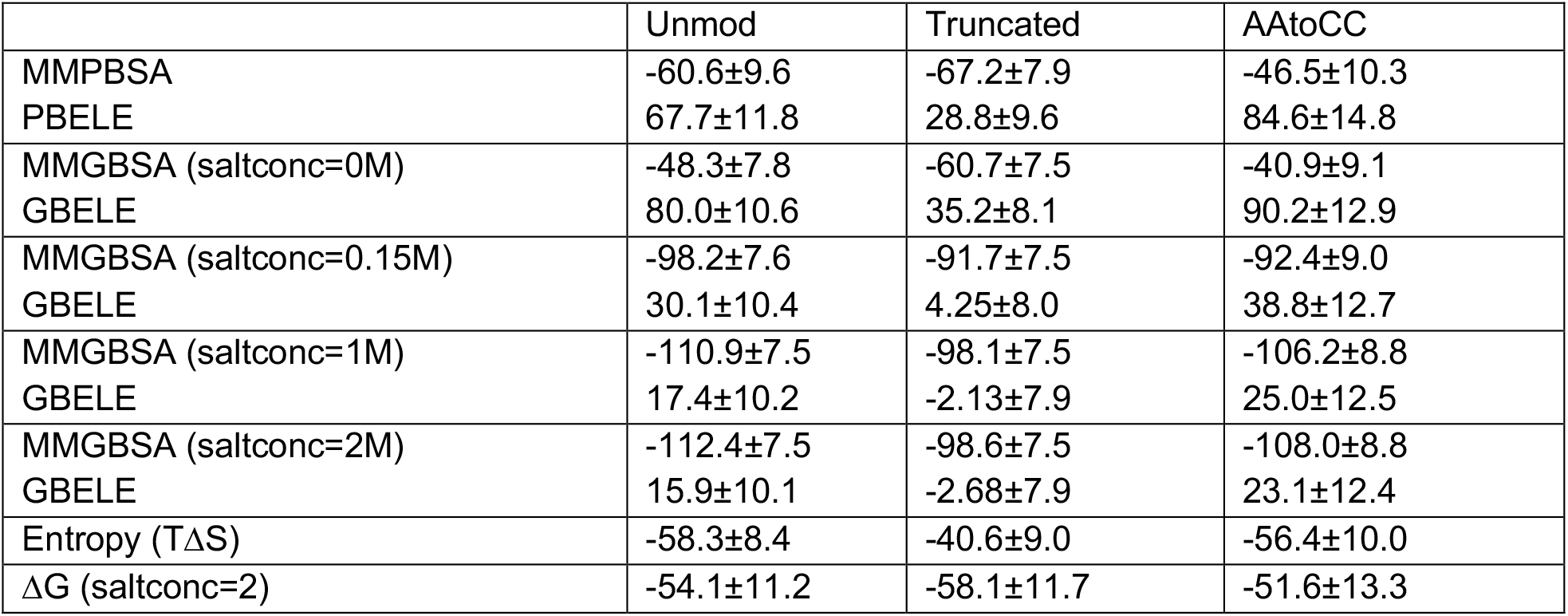
Entropy and free energy of binding analysis as derived by the MMPBSA/MMGBSA analysis.

|  | Unmod | Truncated | AAtCC |
| --- | --- | --- | --- |
| MMPBSA | -60.6±9.6 | -67.2±7.9 | -46.5±10.3 |
| PBELE | 67.7±11.8 | 28.8±9.6 | 84.6±14.8 |
| MMGBSA (saltconc=0M) | -48.3±7.8 | -60.7±7.5 | -40.9±9.1 |
| GBELE | 80.0±10.6 | 35.2±8.1 | 90.2±12.9 |
| MMGBSA (saltconc=0.15M) | -98.2±7.6 | -91.7±7.5 | -92.4±9.0 |
| GBELE | 30.1±10.4 | 4.25±8.0 | 38.8±12.7 |
| MMGBSA (saltconc=1M) | -110.9±7.5 | -98.1±7.5 | -106.2±8.8 |
| GBELE | 17.4±10.2 | -2.13±7.9 | 25.0±12.5 |
| MMGBSA (saltconc=2M) | -112.4±7.5 | -98.6±7.5 | -108.0±8.8 |
| GBELE | 15.9±10.1 | -2.68±7.9 | 23.1±12.4 |
| Entropy (TΔS) | -58.3±8.4 | -40.6±9.0 | -56.4±10.0 |
| ΔG (saltconc=2) | -54.1±11.2 | -58.1±11.7 | -51.6±13.3 |

To compensate for the extensive electrostatic repulsion, we recalculated the MMGBSA energies with higher salt concentrations, which provides more favorable MMGBSA energies. This also changes the binding trend to the same order as the GROMACS energies. However, with the MMGBSA method, the binding energies for all three complexes exhibit a greater degree of similarity compared to the GROMACS interactions energies. The energies begin to converge when the salt concentration reaches 2M, but for the unmodified and the AAtoCC systems, the electrostatic term remains repulsive; suggesting that the energies are underestimated.

We also estimated the entropies for the three systems, as the complex formation is influenced not only by enthalpy but also entropy. We calculated the entropies through normal mode analysis, acknowledging its limitations in terms of accuracy and computational cost.(33) Our calculations show that enthalpy exerts a stronger influence on complex formation. Nevertheless, the entropy term is only 40-50% smaller than the enthalpy term, thus, entropy plays a relatively significant role in the formation of the ASO-RNA hairpin complexes. Although, the entropies overlap within standard deviation, the trend appears reasonable. The truncated ASO system has the least negative entropy term, which can be explained by fewer contacts allowing the RNA hairpin greater freedom of movement in conformational space. The unmodified and AAtoCC systems have similar entropy terms, with 2 kcal/mol less negative for the AAtoCC system; this can be attributed to the greater flexibility of the ASO (Figure 2A-B). Combining the enthalpy and entropy terms, we derive the Gibbs free energy (ΔG). As illustrated in Table 2, the truncated ASO has the most negative ΔG, however, the differences are small and overlap within standard deviations. Thus, with the available computational methods, we cannot successfully reproduce the trend of the experimental determined dissociation constants.

Despite the limitations, valuable insights emerge from the MMGBSA analysis when we dissect the binding energies into interactions between pairs of residues (Figure 3). Firstly, the CG pairs exhibit stronger interaction energies compared to AU pairs, because of more hydrogen bonds, but this does not necessarily imply that these interactions are the determining factor for the stability of the complex. Secondly, in the AAtoCC system, we observe more interactions between ASO bases 6-10 and the RNA 3’-hairpin bases compared to the unmodified system. Furthermore, base 10 of the ASO forms additional interactions with the 5’-terminal bases of the hairpin stem. Although, there are more interactions, the interaction energies are more distributed among several bases and are less attractive, in contrast to the unmodified system. This unsettlement of interactions aligns with the increased flexibility noted in the RMSF analysis for the ASO tail of the AAtoCC system. Thirdly, in the truncated ASO system, there is a decrease in interactions between ASO bases 4-5 and the hairpin, accompanied by an expanded range of interactions involving ASO bases 6-7 with the hairpin compared to the unmodified system. These observations suggest alterations in the dynamics of the complex, which agree with the RMSF profile that shows increased flexibility of the RNA hairpin 3’-stem. For the unmodified system, the ASO base 10 exhibits stronger interactions with base 36 of the 3’-hairpin stem compared to the AAtoCC system. These AU interactions appear to be an important anchoring point, both to stabilize the 3’-tail of ASO and the RNA 3’-stem.

**Figure 3:**
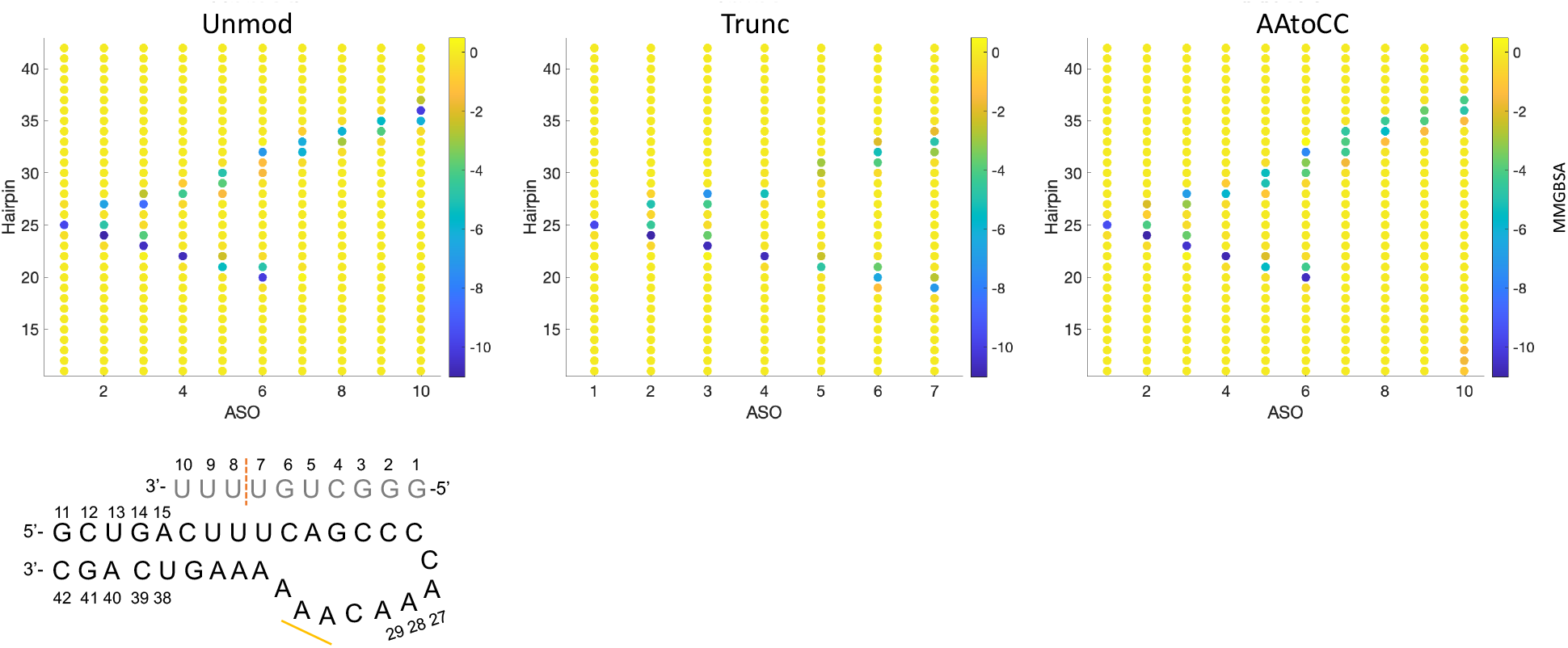
Free energy (from the MMGBSA analysis, in kcal/mol) between ASO-hairpin residue pairs. The schematic figure shows the ASO-hairpin residues numbers and marks the ASO truncation with orange dashed line and the RNA hairpin mutations with yellow bold line.

### Contacts Network and Conformational Analyses

We continue with contacts network analysis and principal component analysis (PCA) to identify key-interactions, confirm the significance of common triple base pairs (b.p.) interactions, and comprehensively map the mechanism underlying the ASO-RNA hairpin complexation. For contacts network analysis we employ our dynamic contacts map approach,(28, 29) where we monitor the evolution of the most frequently occurring specific and nonspecific contacts (present >10% of the time) between pairs of residues in interacting molecules. Specific contacts refer to contacts between the nucleobases, whereas nonspecific – refer to contacts that involve at least one backbone atom (see Methods for details). We follow both the ASO-hairpin contacts and the intramolecular RNA hairpin contacts (Figures 4-5 and S5-S7).

**Figure 4:**
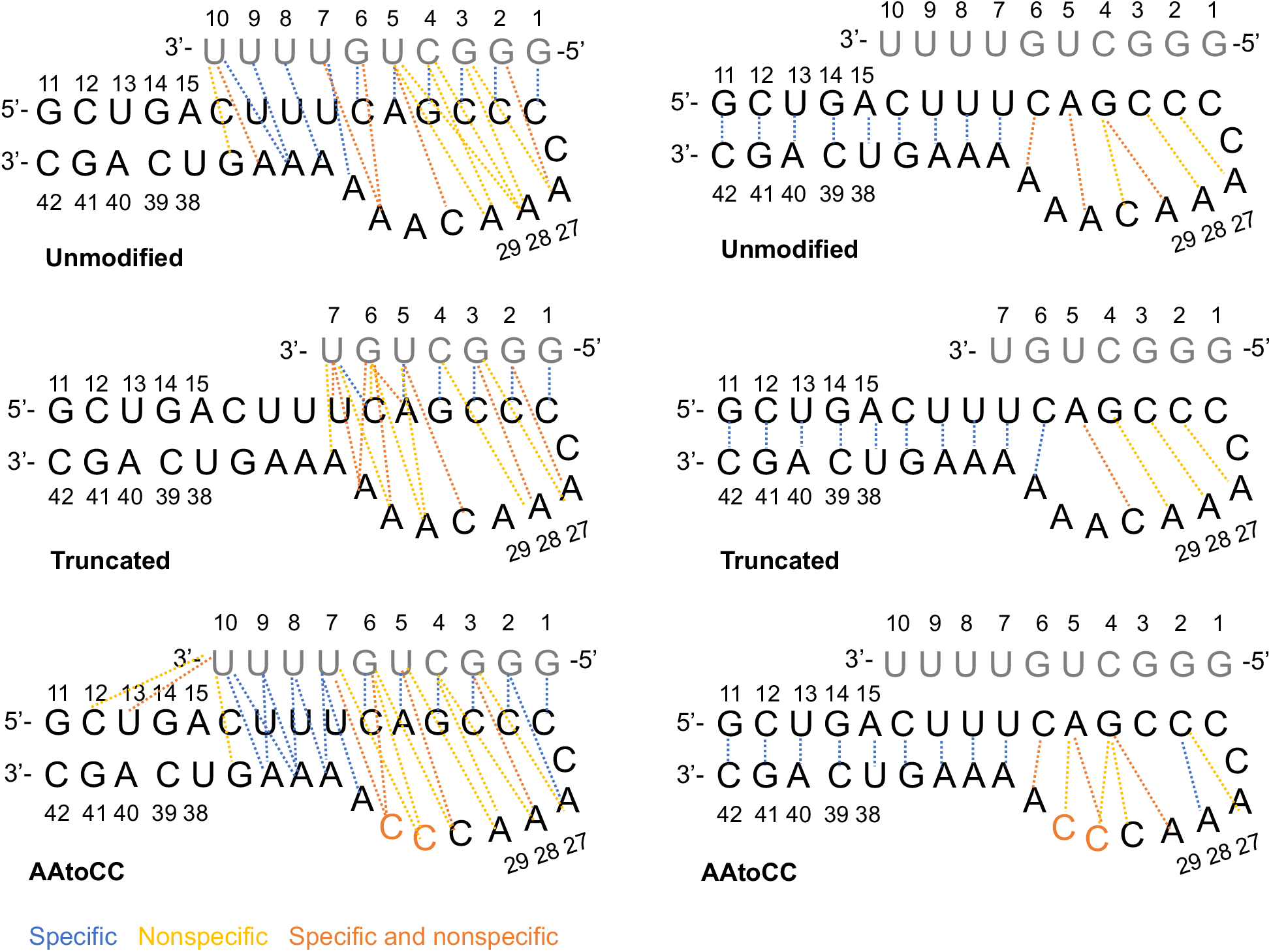
ASO-RNA hairpin interactions and RNA hairpin intermolecular interactions observed in the studied systems.

**Figure 5:**
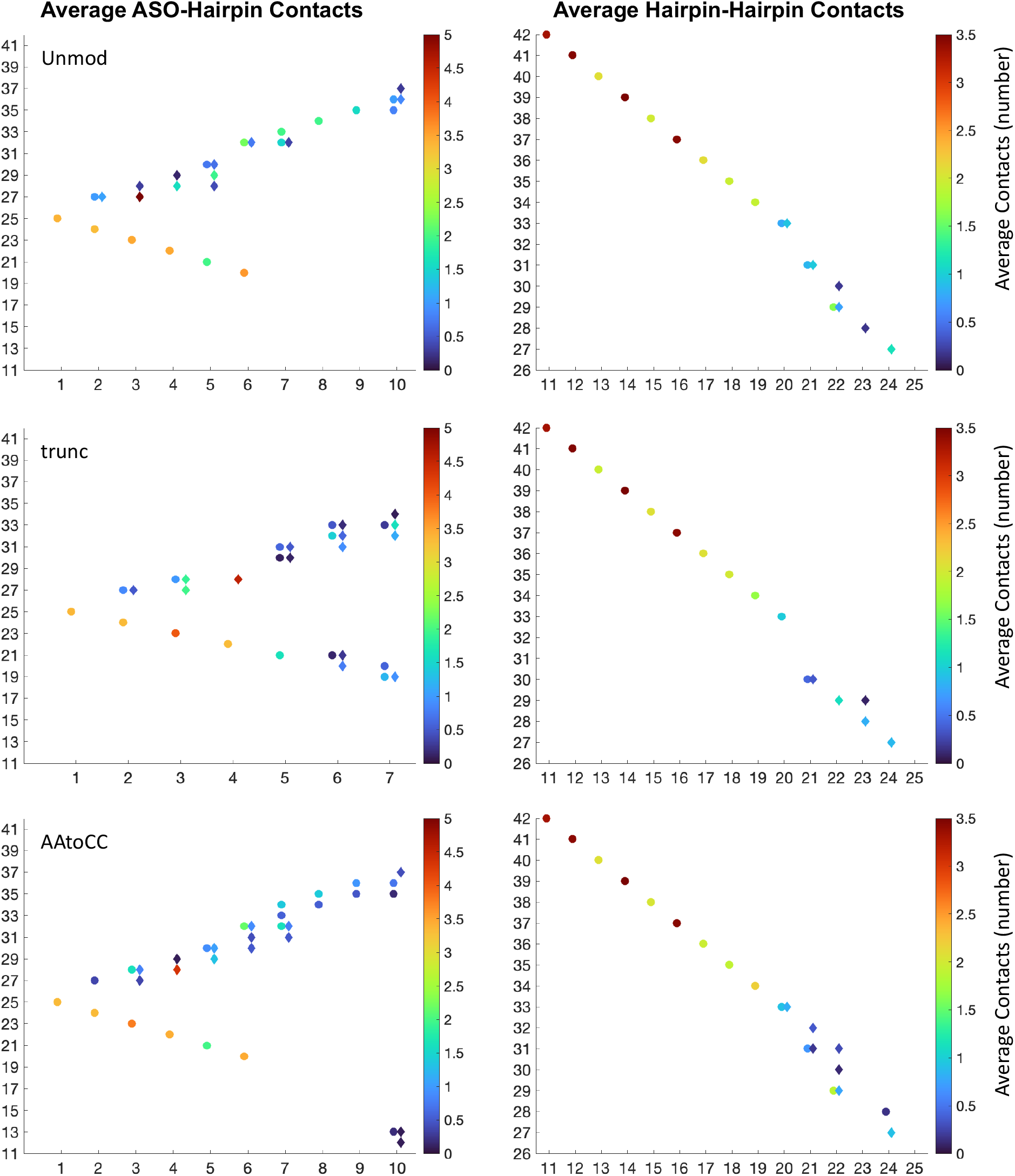
Average contact strength of ASO-hairpin (left-hand panel) and intramolecular RNA hairpin pairs of residues contacts. Specific contacts are denoted with circles and nonspecific contacts are denoted with diamonds.

Analyzing the intramolecular contacts of the RNA hairpin, we observe stable interactions within the stem (bases 11-19, 34-42) across all three systems. Notably, the contacts network within the hairpin loop fluctuates more and vary between the systems, with a greater number of contacts observed for the AAtoCC system, and fewer contacts for the unmodified and truncated ASO systems. For the unmodified system, the loop contacts rearrange at 1500ns (Figure S5), where the G22-A29 contacts transition to G22-C30 and C23-A28 contacts; this agrees with increased RMSF of the A28 and A29 bases. The AAtoCC system shows a similar contacts network as the unmodified system (Figures 4-5), except the G22-A29 interactions, which transition to G22-C30 interactions toward the end (Figure S7). The A31A32->C31C32 mutation contributes to additional contacts involving A21 and G22. The presence of the latter contacts suggests a conformational change in the mutated system towards a more closed loop. The truncated ASO system exhibits a different contacts network, where A21 interacts with C30 instead of A31 (Figures 4-5), and the only stable contacts within the loop being C23-A28 and C24-A27 (Figure S6). Other loop contacts disappear after 1300 ns, which aligns with the large fluctuations in the radius of gyration.

Analyzing the ASO-hairpin contacts (Figures 4-6 and S5-7), we observe a greater number of contacts towards ASO nucleobases 6-10 (AAtoCC system) and 6-7 (truncated system), respectively, compared to the unmodified system. This implies more dynamic and less settled interactions within these two complexes. Contrary, in the unmodified system, the ASO 3’-tail (bases 7-10), maintains consistent interactions with the RNA hairpin stem throughout the trajectory (Figures 1B and 6A, Movie S1). The ITC experiments(12) suggested the interactions formed by the ASO U5 and G6 bases are important for the complex stability. For interactions involving U5 and G6 we observe differences compared to the initial state (Figures 1B and 6A, Movie S2). In the initial state, U5 forms hydrogen bonds with the A21 and A31 bases, and G6 forms hydrogen bonds with the C20 and A32 bases of the hairpin. During MD, the triple b.p. interactions of G6 remain stable. For U5 we observe changes due to the system transition to a more compact state, namely the exchange of the U5-A31 interactions for the A21-A31 interactions. U5 forms also additional contacts with A29 and C30, which later transition to interactions with A28. This is a result of the hairpin bases 28-30 being highly flexible, which through a conformational change, i.e. A29 and C30 base fraying, alter the contacts network at 1500 ns (Figure S4), which explains the change in non-specific interaction energies (Figure S3).

Analyzing the ASO-hairpin contacts for the truncated ASO system (Figures 4, 6B, and S5), we conclude that the missing ASO tail (U8-U10) is crucial for the complex stability. The RNA hairpin becomes more dynamic, resulting in a greater variation of the ASO-hairpin contacts exploited by bases 5-7 (Figure 6B and Movie S3). Initially, we observe that U5 maintains the triple b.p. interactions with A21 and A31 as in the initial state (Figure 1B), G6 only forms nonspecific interactions with C20 and A31 and combined specific and nonspecific interactions with A32, and U7 engages in triple b.p. interactions with C20 and A33. However, at 1500 ns an expansion of the hairpin loop and fraying of C20 occurs, which rearranges the contacts network of U5, G6 and U7. U5 loses its interactions with A31 and reduces its interactions with A21. G6 switches to stacking interactions with A21 and specific interactions with A33. U7 switches to stacking interactions with U19. Finally, we see narrowing of the hairpin loop, recovering the U5 interactions with A21 and A31. G6 forms a new triple b.p. with C20 and A33, and U7 forms a triple b.p. with U19 and A34. Observed fluctuations of the intermolecular contacts suggest an increased likelihood of the complex dissociation. Analyzing the ASO-hairpin contacts for the AAtoCC system (Figure 4 & 6C, S7), we observe that the smaller size of cytosine residues, induce a more compact system, with a dynamically changing contacts network compared to the unmodified system. (Figure 6C, Movie S4). While U5 maintains similar contacts with A21 and C30 as seen for the unmodified system, G6 exhibits different interactions. Initially G6 forms triple b.p. interactions with C20 and C32. However, we observe that G-C-A triple b.p. interactions are more favorable, causing A33 to approach G6 and C20, leading to increased fluctuations of the ASO tail (bases 7-10). U7-U9 shift their interactions from A33-A35 to A34-A36 (Figure 6C and MovieS5). Consequently, U10 becomes more dynamic, engaging in interactions with various bases of the hairpin stem. We believe that the hairpin loop compactness and the enhanced ASO 3’-tail flexibility contribute to the increased repulsion and increased dissociation rate.

**Figure 6:**
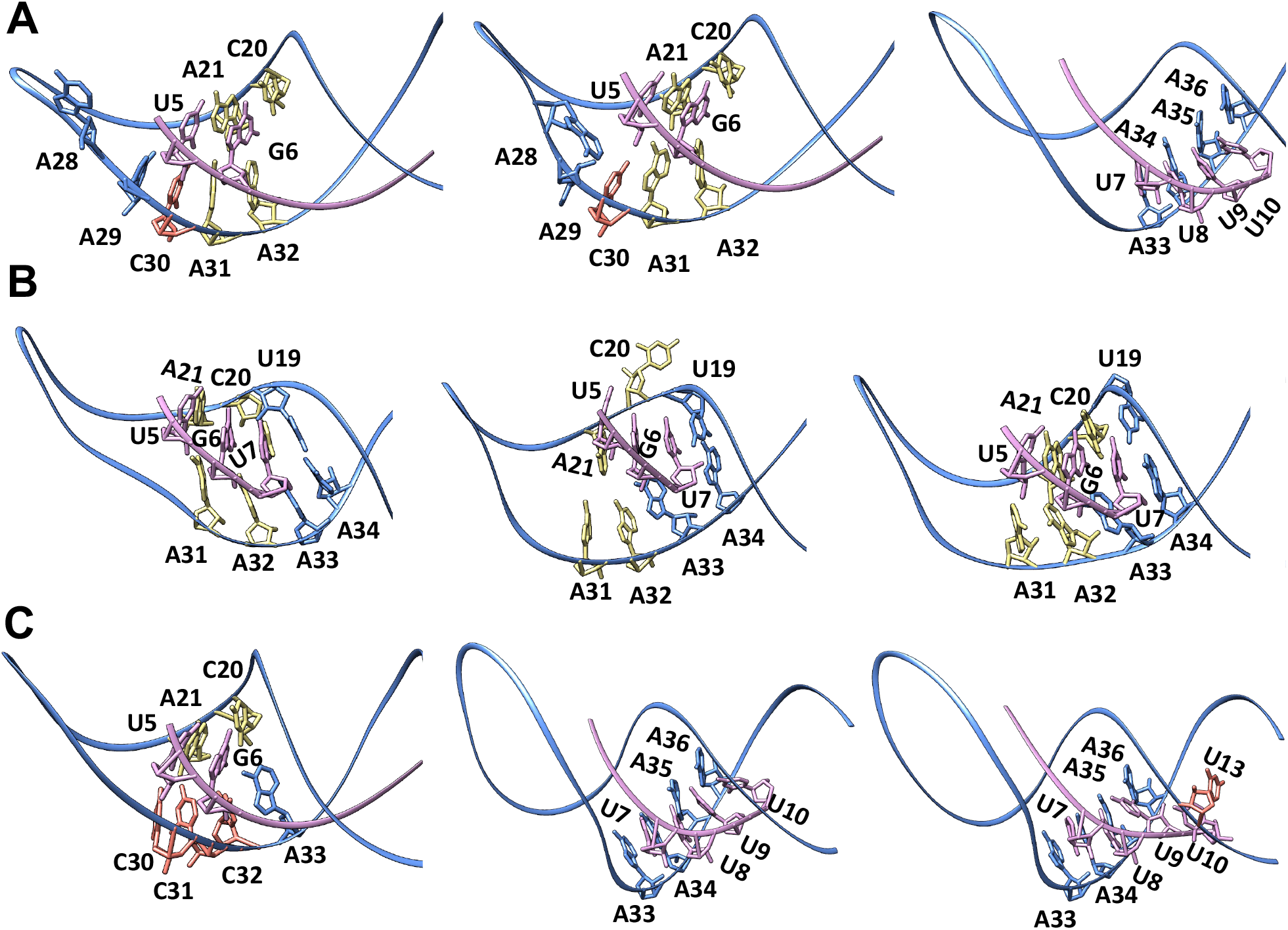
ASO-hairpin contacts seen during MD simulations for the **A**. unmodified, **B**. truncated ASO and **C**. AAtoCC systems.

We also conducted principal component analysis (PCA) based on the complexes RMSD. The first three principal components (PCs) account for 60%, 71%, and 61% of variance in the dynamics for the unmodified, truncated ASO, and AAtoCC systems, respectively (Figure S8). In the unmodified system, these PCs correspond to A28, A29, and C30 base rearrangements (Movies S6-8). In the truncated ASO system, they relate to hairpin loop expansion/narrowing and C20 fraying (Movies S9-11). In the AAtoCC system, they correspond to the compaction of the hairpin loop and the entire complex, U7-U9 rearrangements, and the increased U10 flexibility (Movies S12-14). These results agree with the contacts analysis.

## Conclusions

Antisense oligonucleotide therapy holds great promise for targeting previously challenging targets, offering potential solutions for both rare and common diseases.(2) It also has the potential to revolutionize personalized medicine.(7) However, despite its promise, achieving the desired levels of efficacy, potency and low toxicity profiles remain challenging, as evidenced by the low number of FDA-approved oligonucleotide drugs, with only 18 approvals since 1998.(3) Current ASO design strategies rely heavily on the principle of full Watson-Crick base complementary with various regions within RNA targets, often overlocking the 3D structures of target RNA molecules.(12) While RNAs exhibit a vast conformational landscape, with the ability to fold into diverse 3D shapes that may affect ASO-RNA hybridization. Therefore, elucidating essential mechanistic insights into ASO interactions with different RNA motifs has the potential to expedite the discovery of novel ASO candidates.

With this study we aimed to outline the mechanistic principles governing the stable and selective binding of ASOs to RNA targets. Inspired by Li et al. findings(12), which indicated an advantage of considering common triple base pairs for design of ASOs targeting RNA hairpin motifs, we perform all-atomistic molecular dynamics simulations. Using an RNA pseudoknot modified into an ASO-hairpin complex (Figure 1), we investigated both the impact of the ASO truncation and the mutation (AA to CC, Figure 1) within the hairpin loop. We conducted the analyses of the systems dynamics, inter- and intramolecular interactions, and thermodynamics, and provided detailed mechanistic conclusions for experimentally observed increased dissociation rates in the truncated and mutated systems.

First, the AA to CC mutation in the hairpin loop compacted both the hairpin loop and the complex in general due to the smaller size of cytosines. The cytosines showed a higher propensity to form zippering contacts with the bases of the loop. Consequently, it increased the flexibility of the ASO 3’-tail (bases 7-10). Bases 7-9 shifted their interactions one step down the RNA hairpin stem, while base 10 engaged in diverse interactions with the RNA hairpin stem. The more compact structure of the complex suggests an increase in repulsion forces and dissociation constant. Second, the truncated ASO system revealed that the missing nucleotides of the ASO 3’-tail (bases 8-10) are crucial for the stability of the ASO-hairpin complex. The absence of the ASO tail makes the RNA hairpin more flexible, resulting in increased fluctuations within the ASO-RNA hairpin contact network.

Our computational study further highlights that it is disadvantageous to only consider the Watson-Crick sequence complementarity when designing ASOs. While ASOs may be complementary to one “strand” of an RNA hairpin, the inability of the other “strand” to form favorable triple b.p. interactions can significantly reduce overall affinity.(34) Additionally, the length of the ASO sequence plays a critical role; excessively short ASOs fail to effectively restrict the flexibility of the RNA motif, thereby compromising favorable contacts. However, it is worth noting that longer ASOs may introduce a new challenge. They have the potential to adopt 3D structures, which could lead to interactions with RNA-binding proteins, including transcription factors.(30) These interactions may contribute to ASOs toxicity.

In conclusion, we want to emphasize the significance of considering the 3D structure of an RNA target in ASO design. We propose that pseudoknots and triple helix complexes can be instrumental models for gathering data to create training datasets for the development of machine learning tools, facilitating the design of novel ASO candidates.

## Acknowledgements

The authors thank Swedish National Infrastructure for Computing (SNIC) for the generous provision of computing resources.

## Data Availability

Data is available upon request from the corresponding authors.

