## Supplementary information for "Mechanistic Insights into ASO-RNA Complexation: Advancing Antisense Oligonucleotide Design Strategies"

### Mechanistic Insights for Advancing Antisense Oligonucleotide Design Strategies Targeting RNA Hairpin Motifs

**Table S1**. List of FDA-approved antisense oligonucleotide (ASO) drugs.

| **Name** | **Molecular target/tissue** | **Category/length** | **Approval Date** | **Indications** |
|---|---|---|---|---|
| Fomivirsen (Vitravene) | CMV IE-2/eye | ASO/21 mer ps DNA | 1998.08 | Cytomegalovirus Retinitis |
| Mipomersen (Kynamro) | ApoB-100/liver | ASO/20 mer gapmer, PS | 2013.01 | Homozygous Familial Hypercholesterolemia |
| Eleplinsen (Exondlys 51) | Dystrophin protein/muscle | ASO-SSO****/30 mer, DNA PMO* | 2016.09 | Duchenne Muscular Dystrophy |
| Nusinersen (Spinraza) | Survival motor neuron (SMN) protein/CNS** | ASO/18 mer, PS*** | 2016.12 | Spinal Muscular Atrophy |
| Inotersen (Tegsedi) | TTR/liver | ASO/20 mer gapmer, PS | 2018.01 | Hereditary Transthyretin Amyloidosis, Polyneuropathy |
| Golodirsen (Vyondlys 53) | analogous to Eleplinsen (Exondlys 51) but instead skipping exon 53 | ASO-SSO/25 mer DNA PMO | 2019.12 | Duchenne Muscular Dystrophy |
| Volanesorsen (Waylivra) | Apolipoprotein CIII /liver | ASO/20 mer gapmer, PS | 2019 | Familial Chylomicronemia |
| Viltolarsen (Viltepso) | analogous to Eleplinsen (Exondlys 51) but instead skipping exon 53 | ASO-SSO/21 mer DNA PMO | 2020.08 | Duchenne Muscular Dystrophy |
| Casimersen (Amondys 45) | analogous to Eleplinsen (Exondlys 51) but instead skipping exon 45 | ASO-SSO/22 mer DNA PMO | 2021.02 | Duchenne Muscular Dystrophy |

*Phosphorodiamidate morpholino oligomers (PMO)
**Central nervous system
***Phosphorothioate
****Splice-switching oligonucleotide

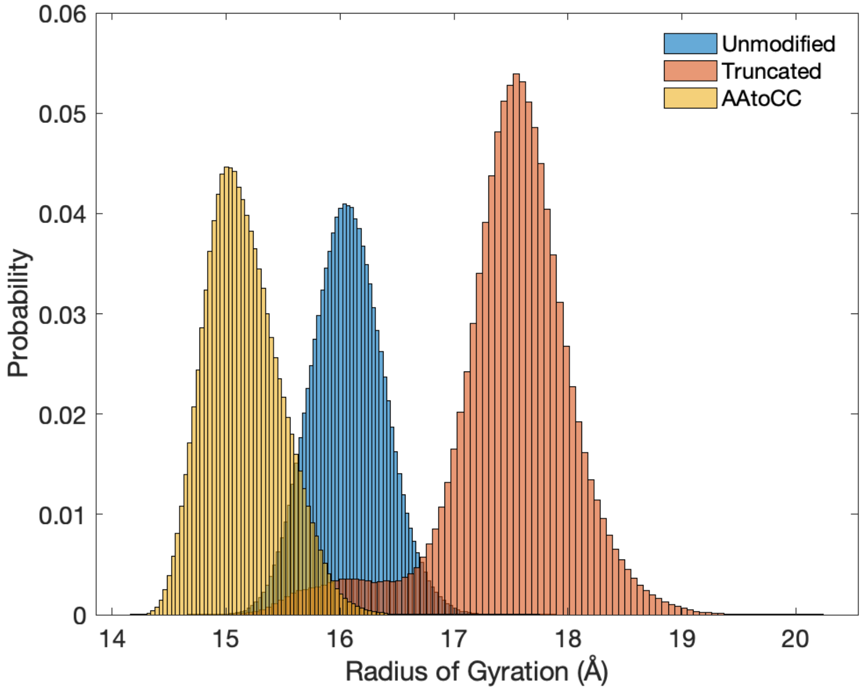


**Figure S1:** Radius of Gyration histogram.

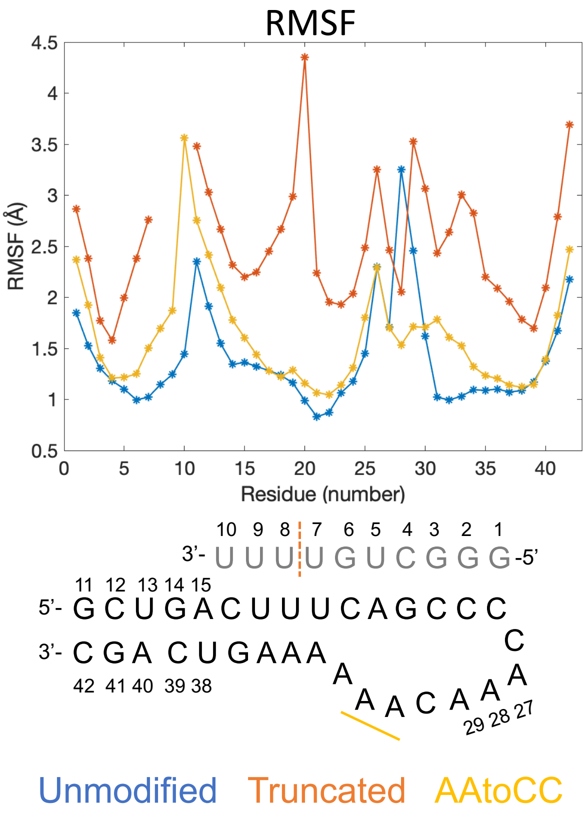


**Figure S2:** Per residue RMSF for the window 100-2000ns. In the schematic figure of the ASO-hairpin, the truncation is marked with orange dashed line and mutation is marked with yellow bold line.

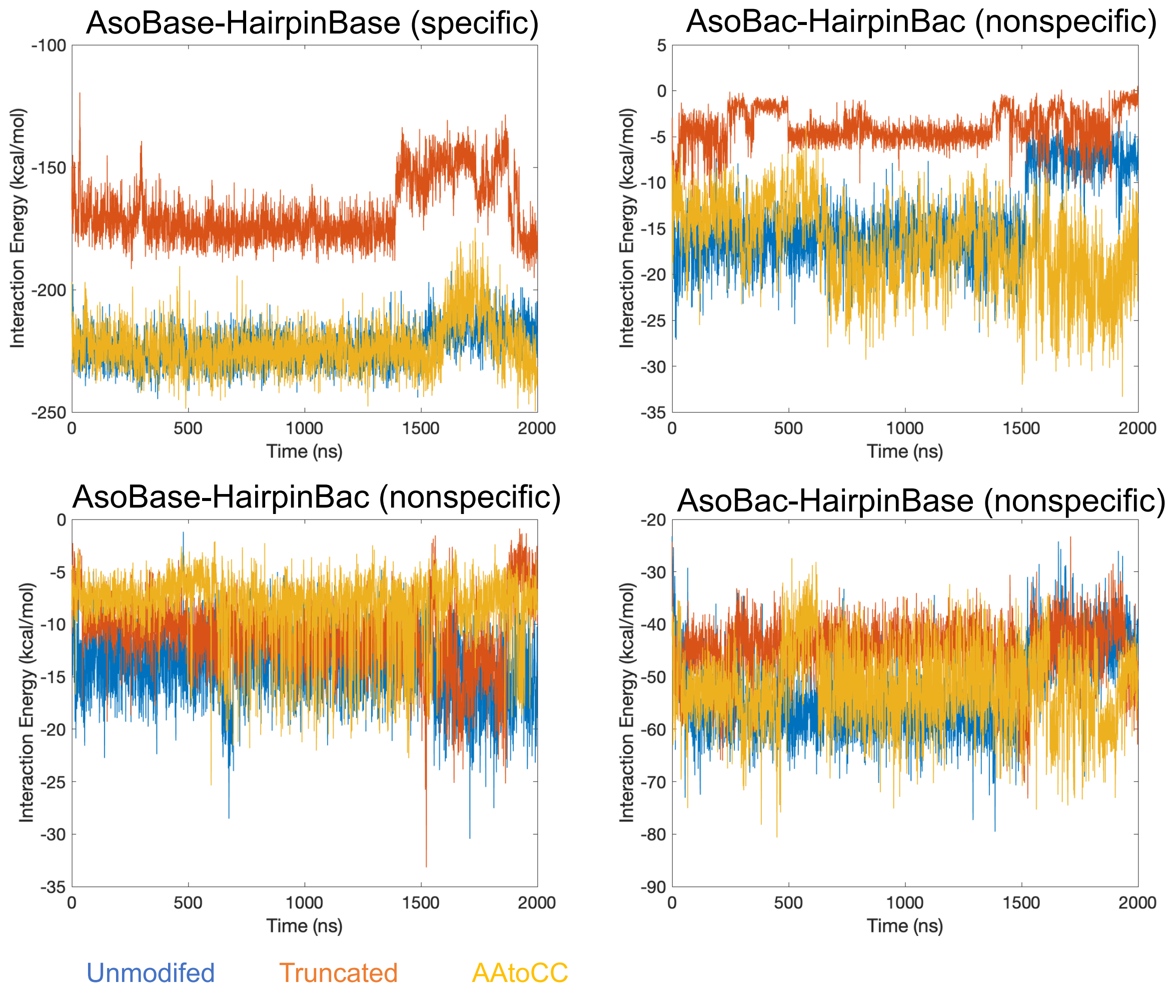


**Figure S3:** GROMACS interaction energies (sum of short-range electrostatic "Coulombic" and vdW "Lennard-Jones" interactions) for different atom groups of ASO-hairpin complex. Base constitutes atoms of the nucleobases and Bac constitutes atoms of the backbone. All interactions involving backbone atoms are referred to as nonspecific interactions.

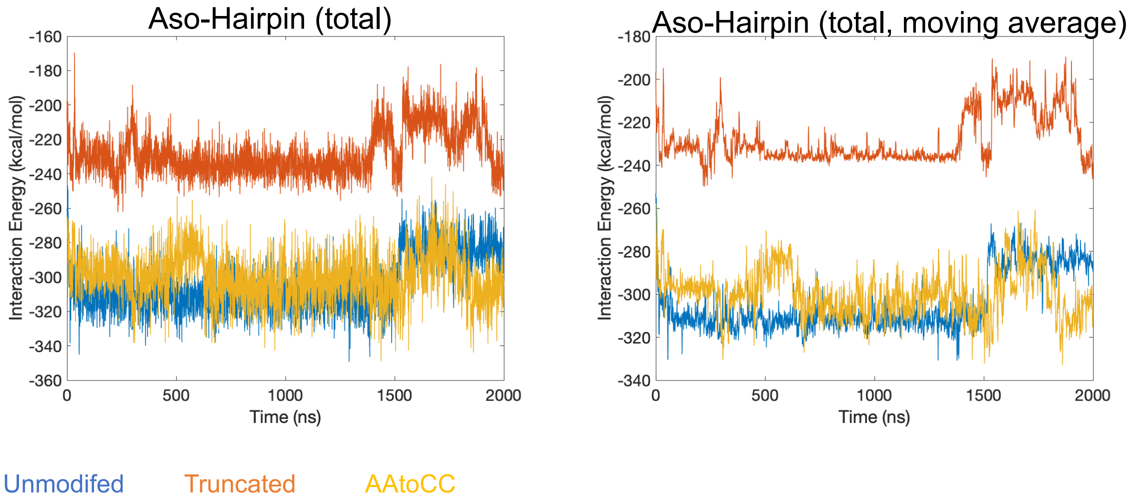


**Figure S4:** GROMACS total interaction energies.

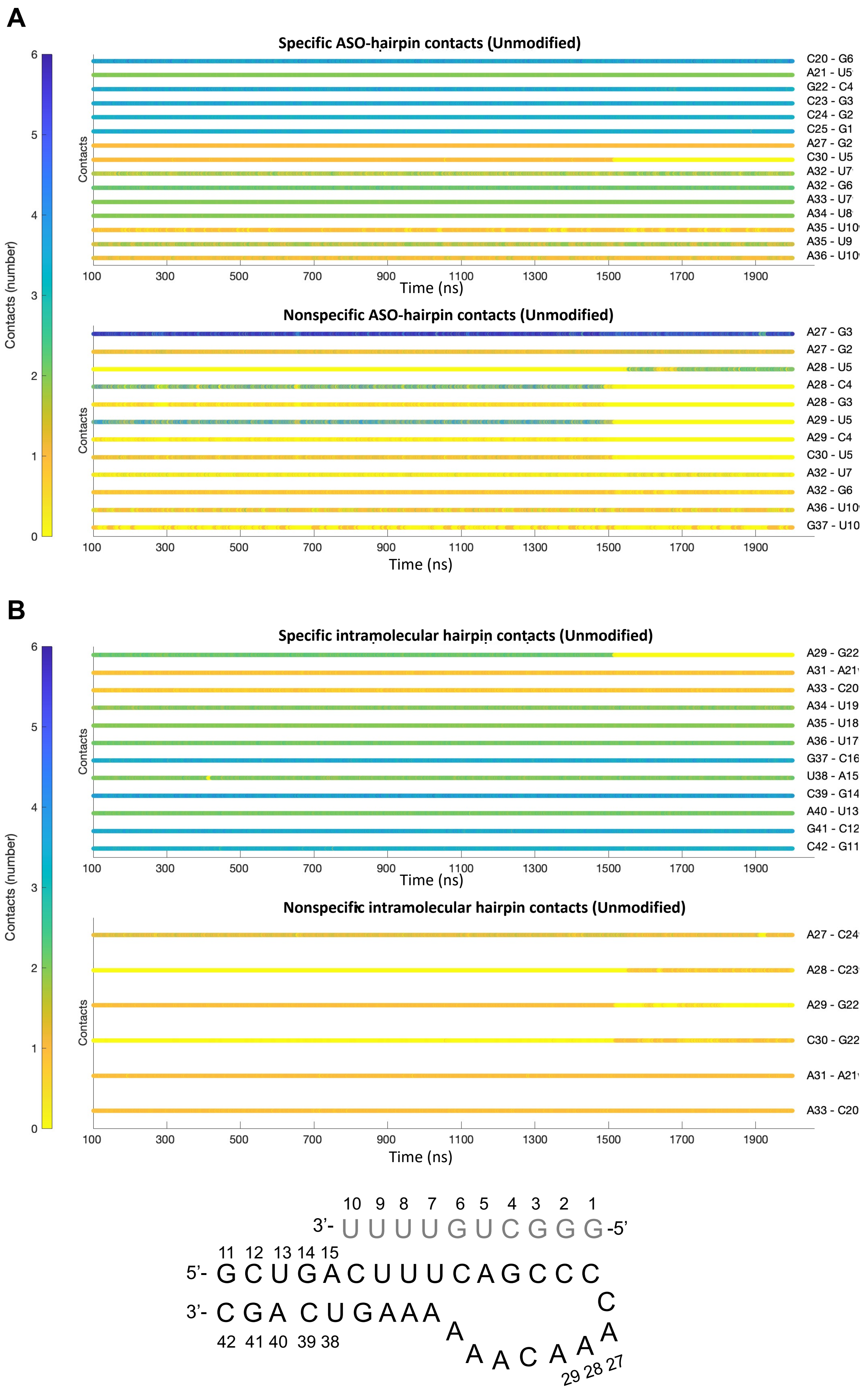
A
Specific ASO-hairpin contacts (Unmodified)
C20 - G6
A21 - U5
G22 - C4
C23 - G3
C24 - G2
C25 - G1
A27 - G2
C30 - U5
A32 - U7
A32 - G6
A33 - U7
A34 - U8
A35 - U10
A35 - U9
A36 - U10
Contacts
Contacts (number)
Time (ns)
Nonspecific ASO-hairpin contacts (Unmodified)
A27 - G3
A27 - G2
A28 - U5
A28 - C4
A28 - G3
A29 - U5
A29 - C4
C30 - U5
A32 - U7
A32 - G6
A36 - U10
G37 - U10
B
Specific intramolecular hairpin contacts (Unmodified)
A29 - G22
A31 - A21
A33 - C20
A34 - U19
A35 - U18
A36 - U17
G37 - C16
U38 - A15
C39 - G14
A40 - U13
G41 - C12
C42 - G11
Nonspecific intramolecular hairpin contacts (Unmodified)
A27 - C24
A28 - C23
A29 - G22
C30 - G22
A31 - A21
A33 - C20
3'- UUUUGUCGGG -5'
5'- GCUGACUUUCAGCCC
3'- CGA CUGAAA A A A C A A A C
42 41 40 39 38
11 12 13 14 15
29 28 27

**Figure S5:** Dynamic contact map for specific and nonspecific **A.** ASO-hairpin contacts and **B.** intramolecular hairpin contacts for the unmodified system.

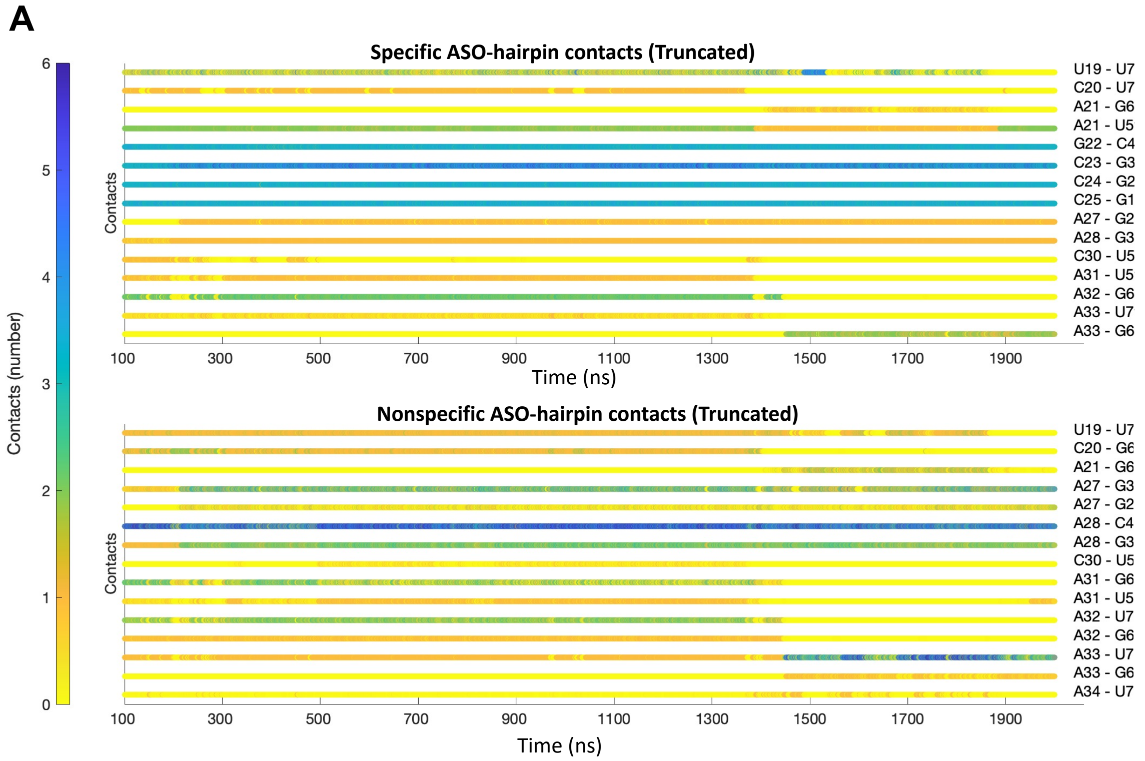

**B**

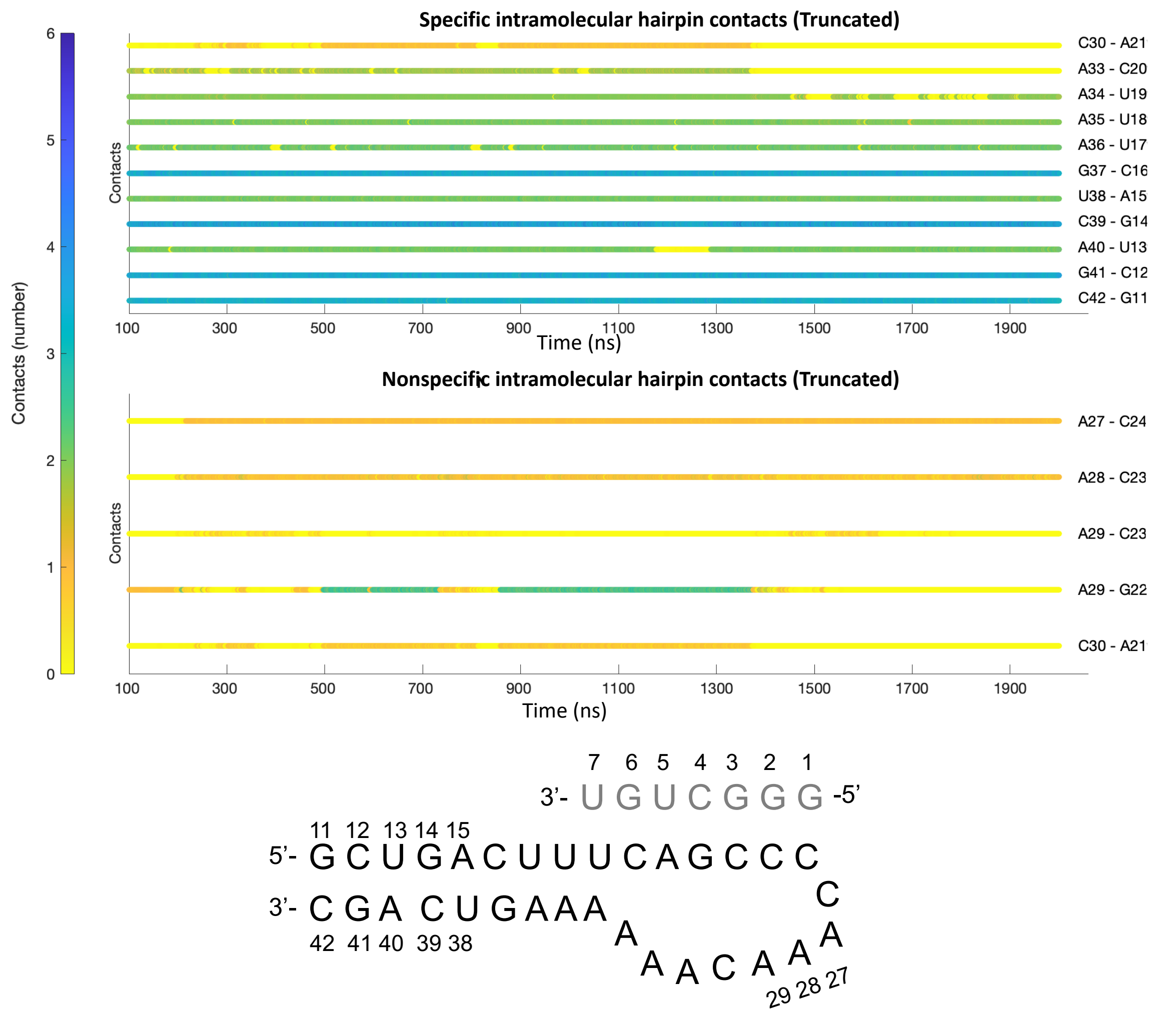


**Figure S6:** Dynamic contact map for specific and nonspecific **A.** ASO-hairpin contacts and **B.** intramolecular hairpin contacts for the truncated ASO system

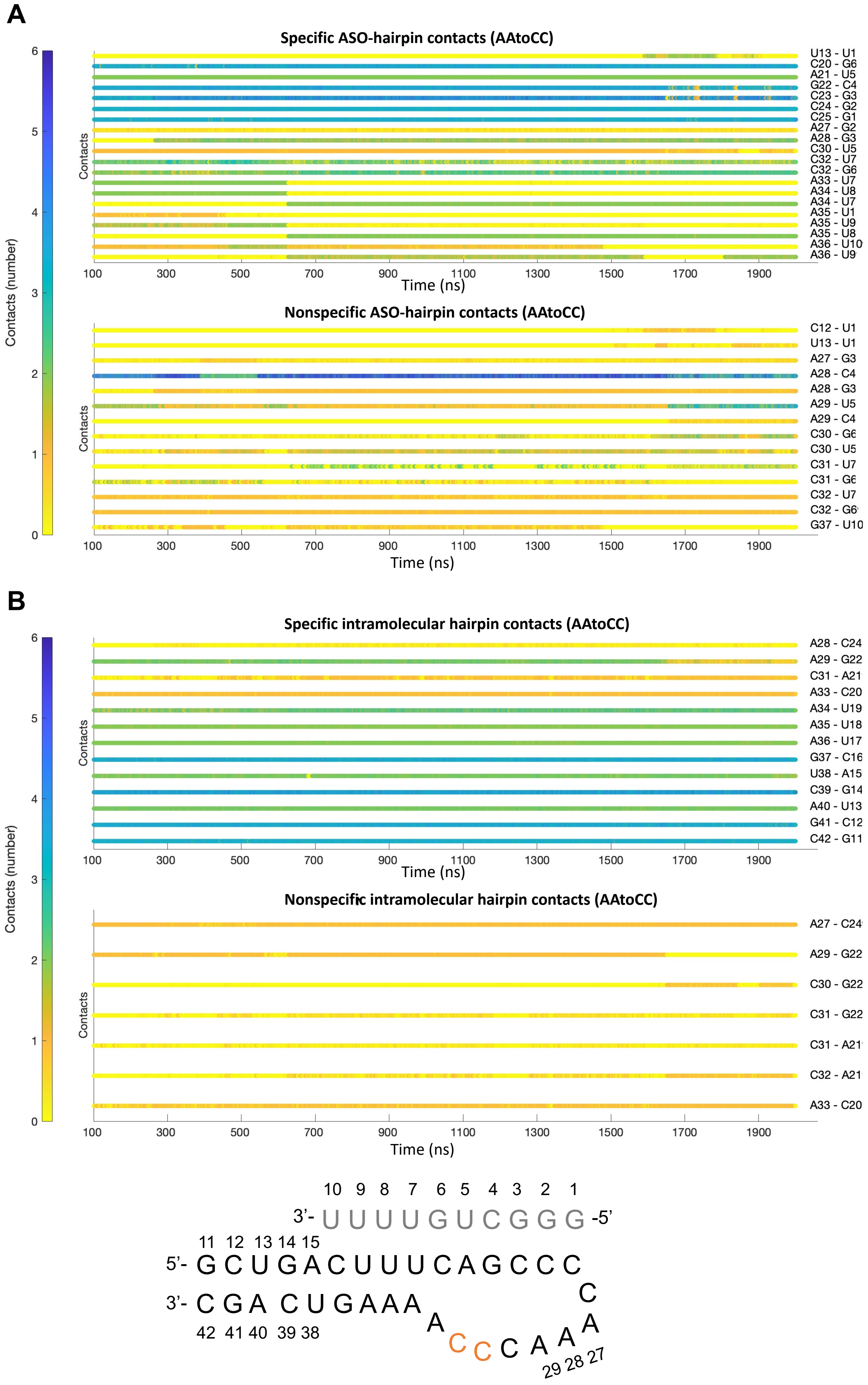
A
Specific ASO-hairpin contacts (AAtoCC)
U13 - U1
C20 - G6
A21 - U5
G22 - C4
C23 - G3
C24 - G2
C25 - G1
A27 - G2
A28 - G3
C30 - U5
C32 - U7
C32 - G6
A33 - U7
A34 - U8
A34 - U7
A35 - U1
A35 - U9
A35 - U8
A36 - U10
A36 - U9
Contacts
Time (ns)
Nonspecific ASO-hairpin contacts (AAtoCC)
C12 - U1
U13 - U1
A27 - G3
A28 - C4
A28 - G3
A29 - U5
A29 - C4
C30 - G6
C30 - U5
C31 - U7
C31 - G6
C32 - U7
C32 - G6
G37 - U10
Contacts
Time (ns)
Contacts (number)
B
Specific intramolecular hairpin contacts (AAtoCC)
A28 - C24
A29 - G22
C31 - A21
A33 - C20
A34 - U19
A35 - U18
A36 - U17
G37 - C16
U38 - A15
C39 - G14
A40 - U13
G41 - C12
C42 - G11
Contacts
Time (ns)
Nonspecific intramolecular hairpin contacts (AAtoCC)
A27 - C24
A29 - G22
C30 - G22
C31 - G22
C31 - A21
C32 - A21
A33 - C20
Contacts
Time (ns)
Contacts (number)
10 9 8 7 6 5 4 3 2 1
3'- U U U U G U C G G G -5'
11 12 13 14 15
5'- G C U G A C U U U C A G C C C
3'- C G A C U G A A A A C C C A A A C
42 41 40 39 38
29 28 27

**Figure S7:** Dynamic contact map for specific and nonspecific **A.** ASO-hairpin contacts and **B.** intramolecular hairpin contacts for the AAtoCC system

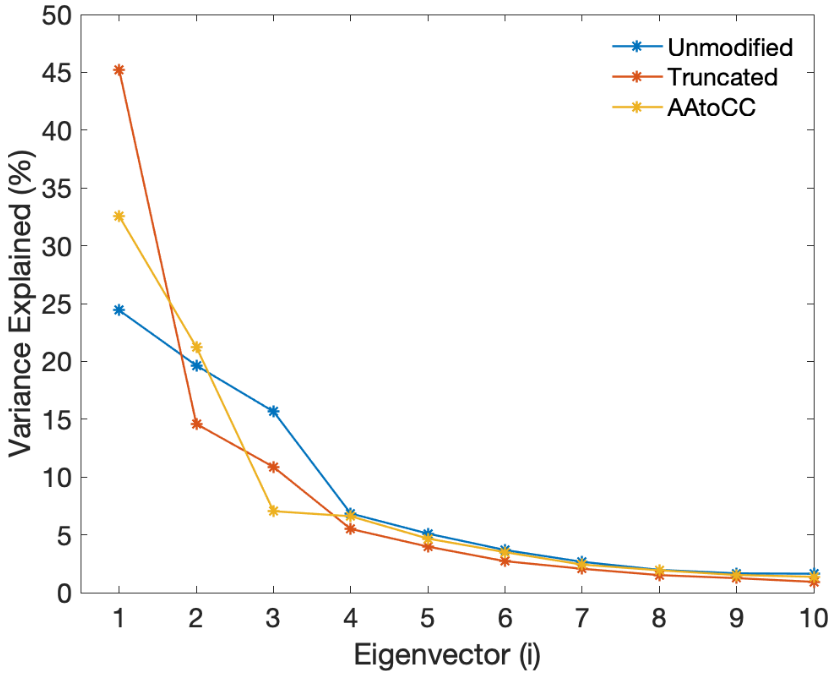


**Figure S8:** Principal component analysis, showing the variance explained for the first 10 eigenvectors.
